# Mathematical modeling explains differential SARS CoV-2 kinetics in lung and nasal passages in remdesivir treated rhesus macaques

**DOI:** 10.1101/2020.06.21.163550

**Authors:** Ashish Goyal, Elizabeth R. Duke, E. Fabian Cardozo-Ojeda, Joshua T. Schiffer

**Author notes:** These authors contributed equally to the work (E. Fabian Cardozo-Ojeda, Joshua T. Schiffer).

## Abstract

Remdesivir was recently demonstrated to decrease recovery time in hospitalized patients with SARS-CoV-2 infection. In rhesus macaques, early initiation of remdesivir therapy prevented pneumonia and lowered viral loads in the lung, but viral loads increased in the nasal passages five days after therapy. We developed mathematical models to explain these results. We identified that 1) drug potency is slightly higher in nasal passages than in lungs, 2) viral load decrease in lungs relative to nasal passages during therapy because of infection-dependent generation of refractory cells in the lung, 3) incomplete drug potency in the lung that decreases viral loads even slightly may allow substantially less lung damage, and 4) increases in nasal viral load may occur due to a slight blunting of peak viral load and subsequent decrease of the intensity of the innate immune response, as well as a lack of refractory cells. We also hypothesize that direct inoculation of the trachea in rhesus macaques may not recapitulate natural infection as lung damage occurs more abruptly in this model than in human infection. We demonstrate with sensitivity analysis that a drug with higher potency could completely suppress viral replication and lower viral loads abruptly in the nasal passages as well as the lung.

**One Sentence Summary:** We developed a mathematical model to explain why remdesivir has a greater antiviral effect on SARS CoV-2 in lung versus nasal passages in rhesus macaques.

## Introduction

There is a desperate need for treatments for SARS CoV-2, the virus which causes COVID-19 disease.^1^ One unmet need for of antiviral therapy development is identification of virologic surrogates for clinically meaningful endpoints such as death or need for hospitalization. In the case of SARS-CoV-2-infected people viral load can be routinely measured in nasal samples or saliva.^2^ However, the primary site of disease is lung tissue. Therefore, bronchoalveolar lavage (BAL) of the lungs would be an ideal sample. However, BAL is usually not necessary for diagnosis, represents an infection risk to medical personnel and is rarely performed in the care of COVID-19 patients. When BAL does occur, it is often late during disease in critically ill patients rather than at early clinical presentation. Thus, the natural kinetics of SARS CoV-2 in lungs are unknown in humans.

In humans, a double-blind, randomized, placebo-controlled trial showed that the nucleoside analog remdesivir limited the duration of illness and approached statistical significance for reduction in mortality when given later in disease^3^. In a separate study with an overall later time of treatment initiation, remdesivir had no effect on viral load or clinical outcome.^4^ A recent pre-print demonstrated that remdesivir was highly effective when initiated 12 hours after infection in rhesus macaques.^5^ In this context, remdesivir prevented pneumonia and limited extent of clinical illness. While there was an effect on viral shedding in serial BAL specimens, viral load in the nasal passage was unchanged relative to animals treated with a vehicle during the first several days of infection and higher starting at day 5. A critical question is whether nasal viral loads are potentially useful as a surrogate for the extent of lung disease during COVID-19 infection.

Here, we develop mathematical models that recapitulate viral load trajectories in both anatomic compartments of the infected rhesus macaques. The models explain these differences according to different underlying viral kinetics off therapy in both compartments as well as differential drug potency in nasal passage versus lung.

## Results

### Discrepant SARS CoV-2 viral loads in lungs and nasal passages in response to remdesivir treatment in rhesus macaques

In a recent pre-print article, 12 rhesus macaques were infected with 2.6×10^6^ TCID50 of SARS-CoV-2 strain nCoV-WA1-2020 via intranasal, oral, ocular and intratracheal routes and then treated with either placebo or IV remdesivir (10 mg/kg loading dose followed by 5 days of 5 mg/kg) starting at 12 hours post infection.^5^ Overall treatment resulted in reduced severity of clinical illness, less pronounced infiltrates on chest radiograph, lower viral load by nucleic acid and viral titer measurement in bronchoalveolar lavage fluid on days 1, 3 and 7, and decreased volume of lung lesions, lung weight and inflammation on histologic post-mortem exam.^5^

We re-examined viral loads in BAL and nasal specimens, and noted that at days 1, 3 and 7 post-infection BAL viral loads were lower in the remdesivir arm relative to the vehicle arm by approximately a single order of magnitude **(Fig 1a)**. Similar results were observed when viral load was measured using viral culture.^5^ In nasal specimens, there was no observed difference in viral loads at days 1, 2, 3 and 4; on day 6, there was a trend towards higher viral loads in the remdesivir treated arm; on day 5 and 7, nasal viral loads were statistically higher in the treated arm relative to vehicle **(Fig. 1b)**.

**Figure 1:**
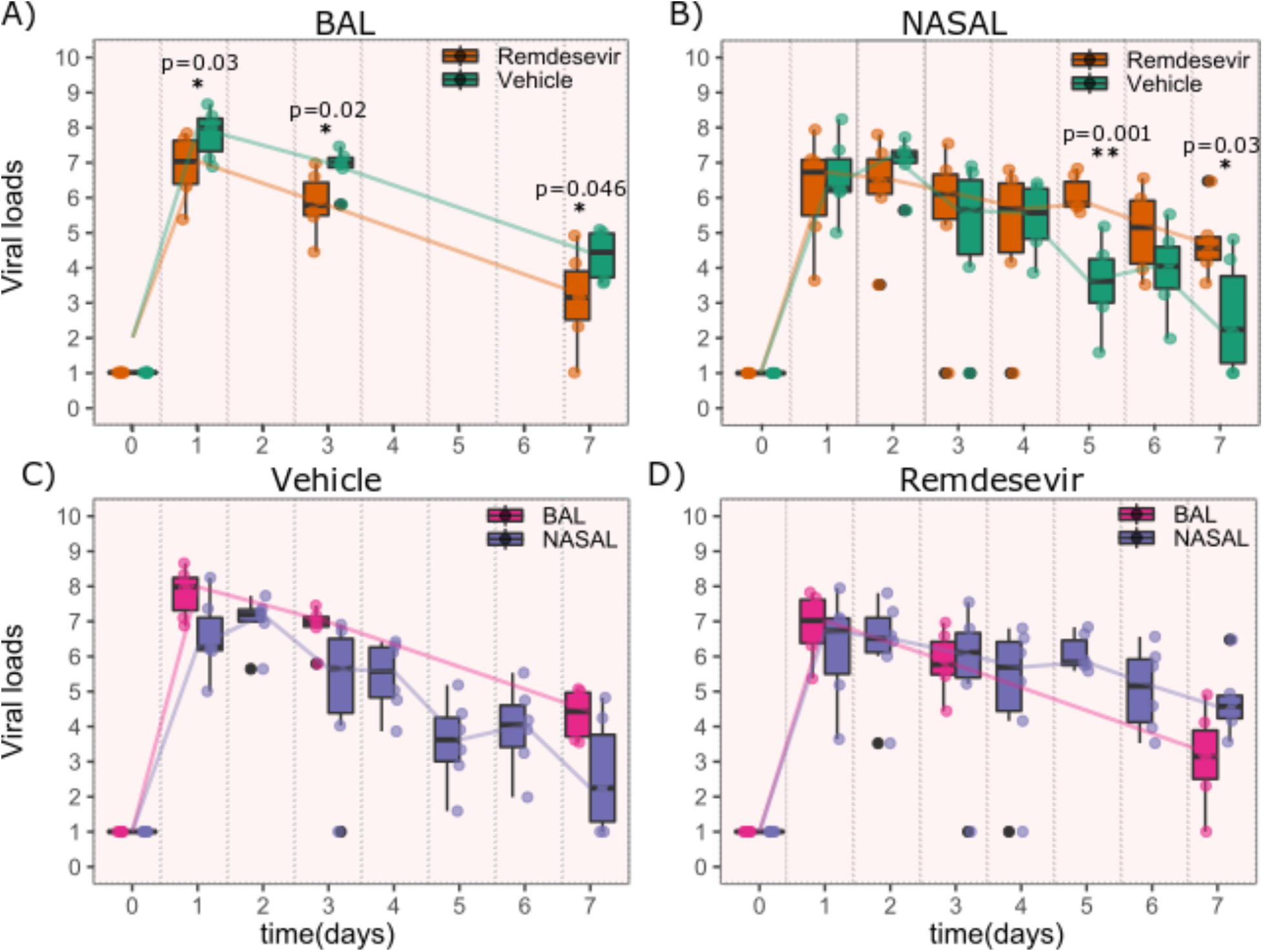
Viral load kinetics following remdesivir treatment in 6 rhesus macaques. **A**. Decreased bronchoalveolar lavage (BAL) viral loads in 6 remdesivir treated animals versus 6 vehicle controls at all measured time points. **B.** Increased nasal viral loads in remdesivir treated animals versus controls at late timepoints. **C**. BAL versus nasal viral loads in vehicle treated animals. **D**. BAL versus nasal viral loads in remdesivir treated animals. The Wilcoxon rank sum test was employed to determine the differences in the median viral loads in treated and vehicle controls at different time points. p<0.05 denotes statistical significance. (*); p < 0.01 is denoted **.

When nasal viral loads were compared longitudinally to BAL viral loads in vehicle treated animals, viral loads were generally higher in BAL than in the nasal passages at days 1, 3 and 7 **(Fig 1c)**; in the remdesivir-treated animals, viral loads were equivalent on days 1 and 3, but higher in nasal passages than BAL at day 7 **(Fig 1d)**. Overall, these results suggest that remdesivir lowered viral load in the lung but appeared to have the opposite effect in nasal passages of rhesus macaques at late timepoints.

### Dual compartment PK/ PD model of remdesivir

To explain the differential observations in lung and nasal passages of remdesivir-treated animals, we developed a mathematical model to capture drug pharmacokinetic (PK) and pharmacodynamics (PD), as well as viral and immune dynamics. The PK model is represented in **Fig 2** with equations listed in the **Methods.** It captures the steps following intravenous injection of remdesivir (GS-5734), including conversion to an alanine metabolite (GS-704277) and then to the parent nucleoside GS-441524 (Nuc), the necessary phosphorylation of this molecule to achieve its active triphosphate form (NTP) as well as the distribution of these metabolites from plasma to tissue.

**Figure 2:**
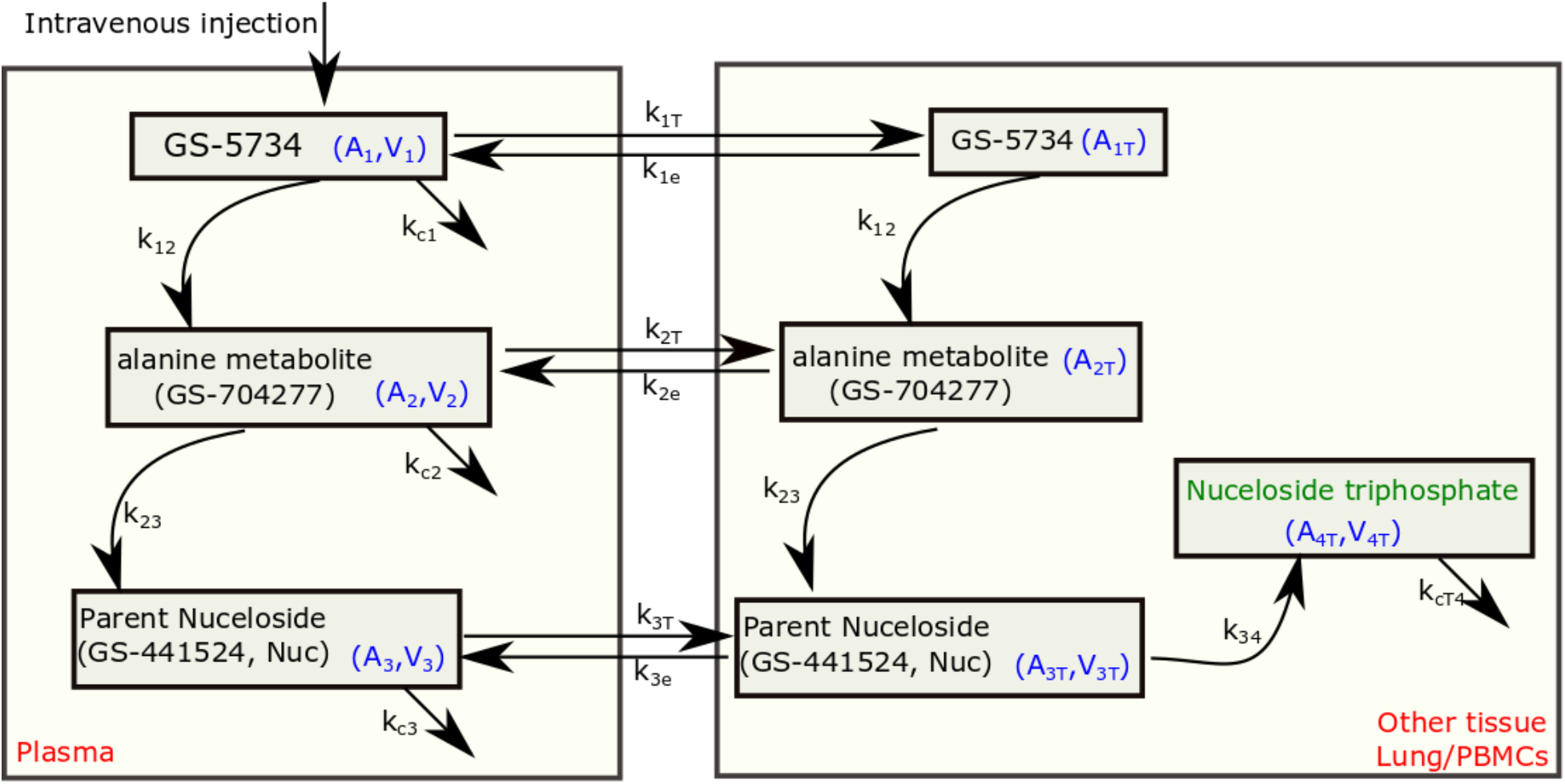
Schematic of the remdesivir pharmacokinetic (PK) model. The model includes plasma and tissue levels of remdesivir GS-5734 (*A*_1_, *A*_1T_), the alanine metabolite GS-507277 (*A*_2_, *A*_2T_) and the parent nucleoside GS-441524 (*A*_3_, *A*_3T_) that is phosphorylated in tissue to the active nucleoside triphosphate form of the drug (*A_4T_*). For remdesivir and the first two metabolites, we modeled the drug distribution from plasma to tissues.

For single-dose PK, we fit the model to data from healthy rhesus macaques in which various intermediate metabolites were measured over time following a single injection of 10 mg/kg, including the levels of NTP is PBMCs.^6,7^ We also simultaneously fit the model to multi-dose drug and metabolite trough levels from the infected rhesus macaques (10 mg/kg at day 0.5 and thereafter, 5 mg/kg daily at days 2 till 6 post-infection), including a day 7 level of the Nuc in tissue at the time of necropsy on day 7 in **Fig 3b.**^5^ RDV PK parameters are listed in **Table 1.** The PD models assumes an EC_50_ or concentration of the active metabolite (NTP), at which replication is inhibited by 50%.

**Figure 3:**
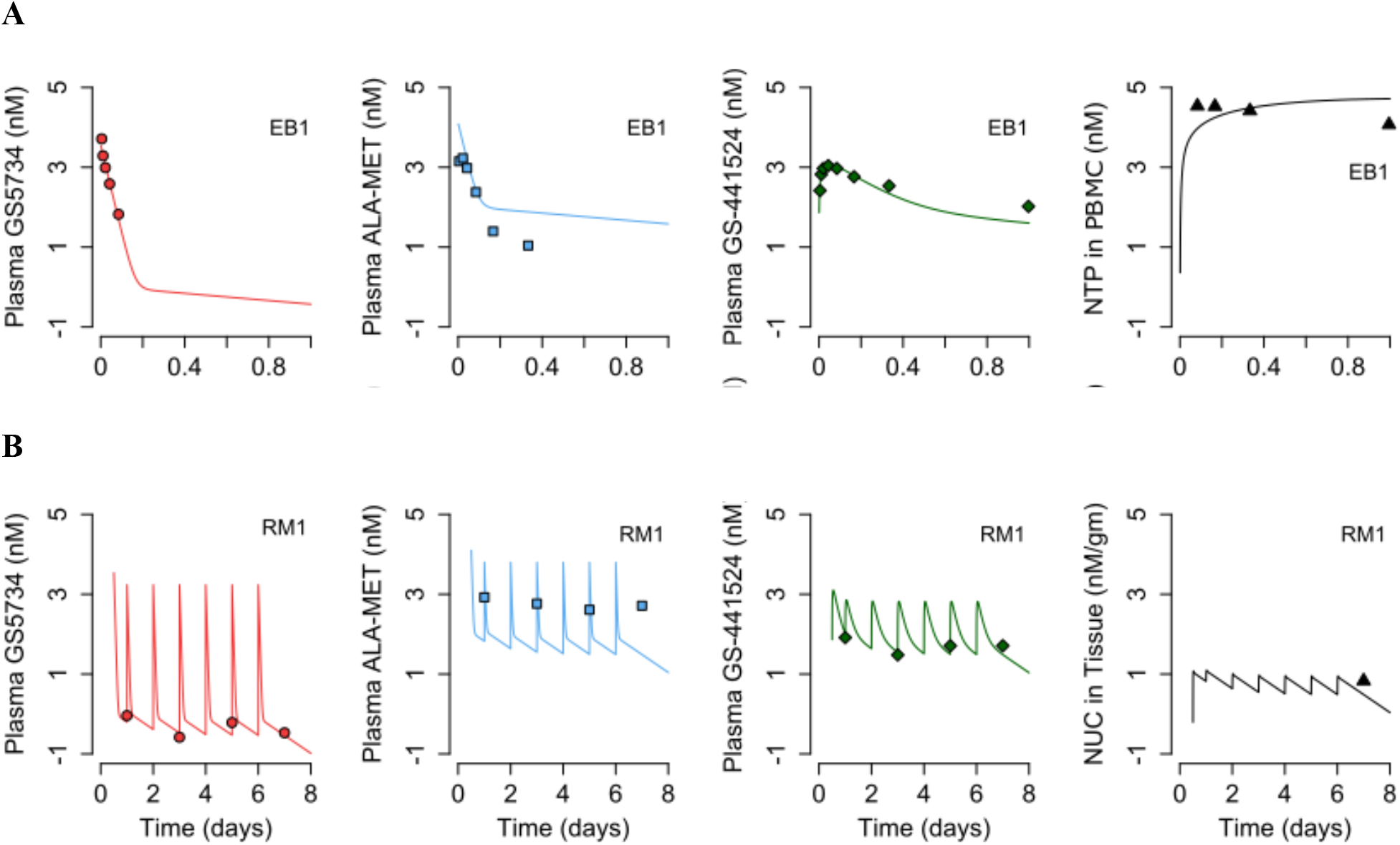
Remdesivir (RDV) pharmacokinetic (PK) model fits to data. **A.** RDV PK model fits to data in rhesus macaques from a single dose of 10mg/kg remdesevir at day 0. **B.** Representative fits of drug and intermediate levels from one of the rhesus macaques (RM1) from the current COVID-19 study. Animals received 10mg/kg remdesevir at day 0.5 and 5mg/kg remdesevir at days 1, 2, 3, 4, 5 and 6. The other animals RM2-RM6 have similar RDV PK profiles.

**Table 1.** Pharmacokinetic model parameters.

| | $k_{1T}$ | $k_{2T}$ | $k_{1e}$ | $k_{2e}$ | $k_{12}$ | $k_{23}$ | $k_{34T}$ | $k_3$ | $V_1$ | $V_2$ | $V_3$ | $V_{3T}$ | $V_{4T}$ |
| --- | --- | --- | --- | --- | --- | --- | --- | --- | --- | --- | --- | --- | --- |
| RM1 | 33.0 | 5.86<br>$\times 10^3$ | 0.02 | 12.3 | 1.0 | 989.6 | 158.4 | 7.9 | $2.8 \times 10^{-3}$ | $1.1 \times 10^{-7}$ | $1.7 \times 10^{-5}$ | $3.5 \times 10^{-3}$ | $5.8 \times 10^{-5}$ |
| RM2 | 33.0 | 5.86<br>$\times 10^3$ | 0.02 | 12.3 | 1.0 | 989.9 | 168.9 | 7.9 | $2.8 \times 10^{-3}$ | $1.1 \times 10^{-7}$ | $1.8 \times 10^{-5}$ | $4.0 \times 10^{-3}$ | $5.8 \times 10^{-5}$ |
| RM3 | 33.0 | 5.86<br>$\times 10^3$ | 0.02 | 12.3 | 1.0 | 989.7 | 162.4 | 7.9 | $2.8 \times 10^{-3}$ | $1.1 \times 10^{-7}$ | $1.7 \times 10^{-5}$ | $3.7 \times 10^{-3}$ | $5.8 \times 10^{-5}$ |
| RM4 | 33.0 | 5.86<br>$\times 10^3$ | 0.02 | 12.3 | 1.0 | 990.1 | 173.4 | 7.9 | $2.8 \times 10^{-3}$ | $1.1 \times 10^{-7}$ | $1.8 \times 10^{-5}$ | $4.2 \times 10^{-3}$ | $5.8 \times 10^{-5}$ |
| RM5 | 33.0 | 5.86<br>$\times 10^3$ | 0.02 | 12.3 | 1.0 | 990.1 | 166.9 | 7.9 | $2.8 \times 10^{-3}$ | $1.1 \times 10^{-7}$ | $1.8 \times 10^{-5}$ | $3.9 \times 10^{-3}$ | $5.8 \times 10^{-5}$ |
| RM6 | 33.0 | 5.86<br>$\times 10^3$ | 0.02 | 12.3 | 1.0 | 989.8 | 175.8 | 7.9 | $2.8 \times 10^{-3}$ | $1.1 \times 10^{-7}$ | $1.8 \times 10^{-5}$ | $4.3 \times 10^{-3}$ | $5.8 \times 10^{-5}$ |

The model is able to recapitulate the levels of remdesivir and its metabolites in healthy (**Fig 3a**) and infected rhesus macaques **(Fig 3b)**.

### Lung and nasal mathematical model of SARS Co-V-2 in rhesus macaques

We developed a model of viral replication in the nasal passage and lungs that includes multiple mechanisms that may occur following infection in lungs and the nasal passages **(Fig 4a**). This model is an adaptation of our previous model of human COVID-19 infection and includes a density-dependent death of infected cells as a proxy for an intensifying innate response to a higher burden of infection, proliferation of susceptible epithelial cells, the conversion of susceptible and/or infected cells to an infection refractory state and the movement of the virus between nasal passages and the lung.^8^ We fit different versions of this model to the viral loads in nasal passages and lung from 6 infected, remdesivir-treated animals,^5^ and 14 infected and untreated animals, including the 6 vehicle animals in the remdesivir protocol (RM7-12),^5^ and 8 other animals infected using the same protocol (4 of which had extended nasal viral load measures through 21 days after infection).^9^ Each model explored was a version of the full model in **Fig 4a** with individual components removed (**Methods**)

**Figure 4:**
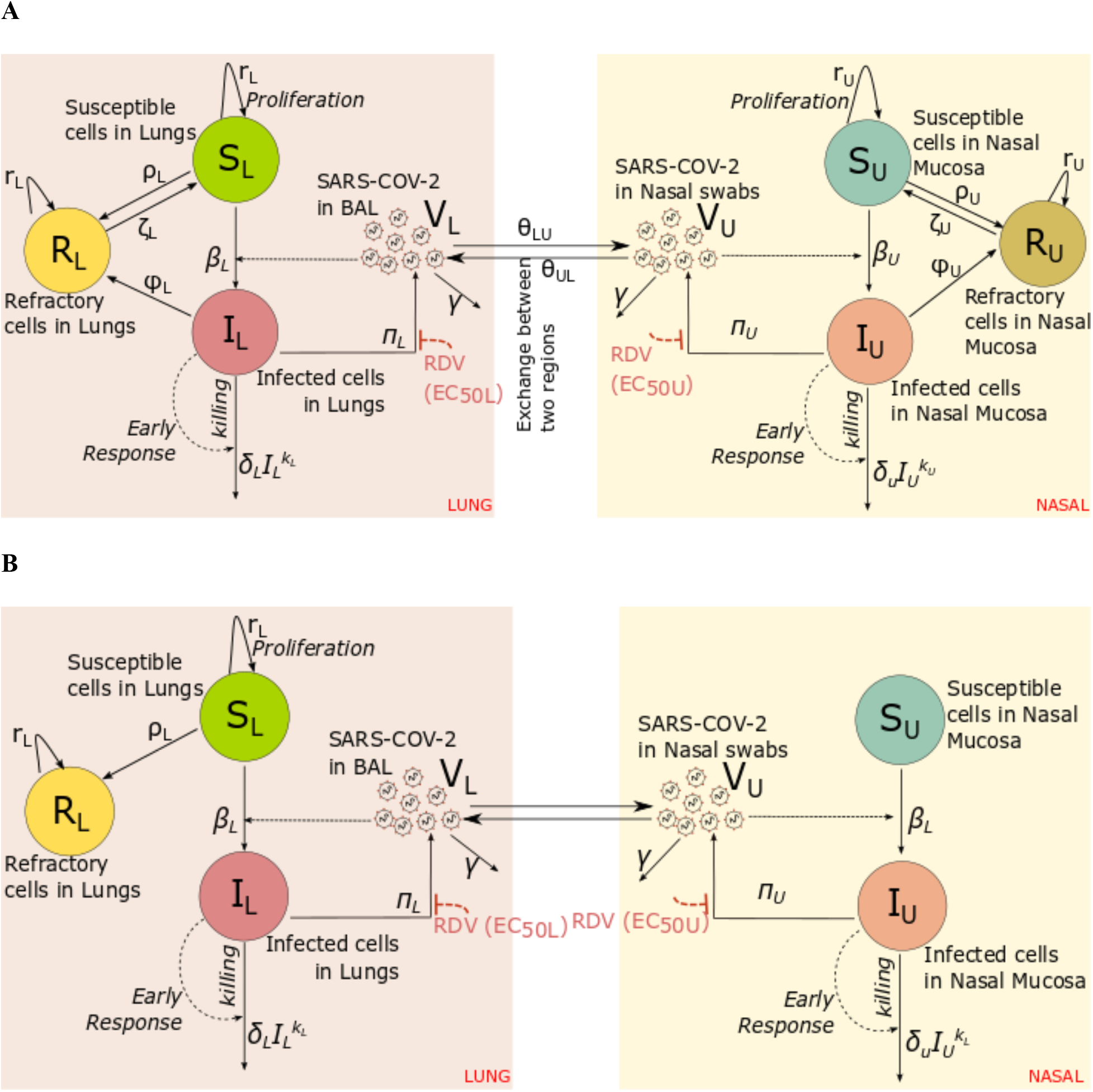
Mathematical models of nasal and lung SARS CoV-2 dynamics and remdesivir therapy. **A.** Schematic of a comprehensive viral dynamics model inclusive of all possible compartments and assumptions. **B.** A reduced model that recapitulates the complete viral load data. Exclusions relative to the complete model include no refractory cell compartment in the nasal passage and no proliferation of susceptible cells in the nasal compartment.

Using model selection theory, we found that the model with minimal complexity necessary to explain the observed data was the one in **Fig. 4b**. In this model, infected cell death and viral production have different rates in lung compared to the nasal passage (**Table 2**). Furthermore, in the selected model, susceptible lung cells proliferate and become refractory to infection, but cells in the nasal passages do not (**Table 3**). Interestingly, this model lacked viral interchange between lung and nasal passages.

**Table 2:**
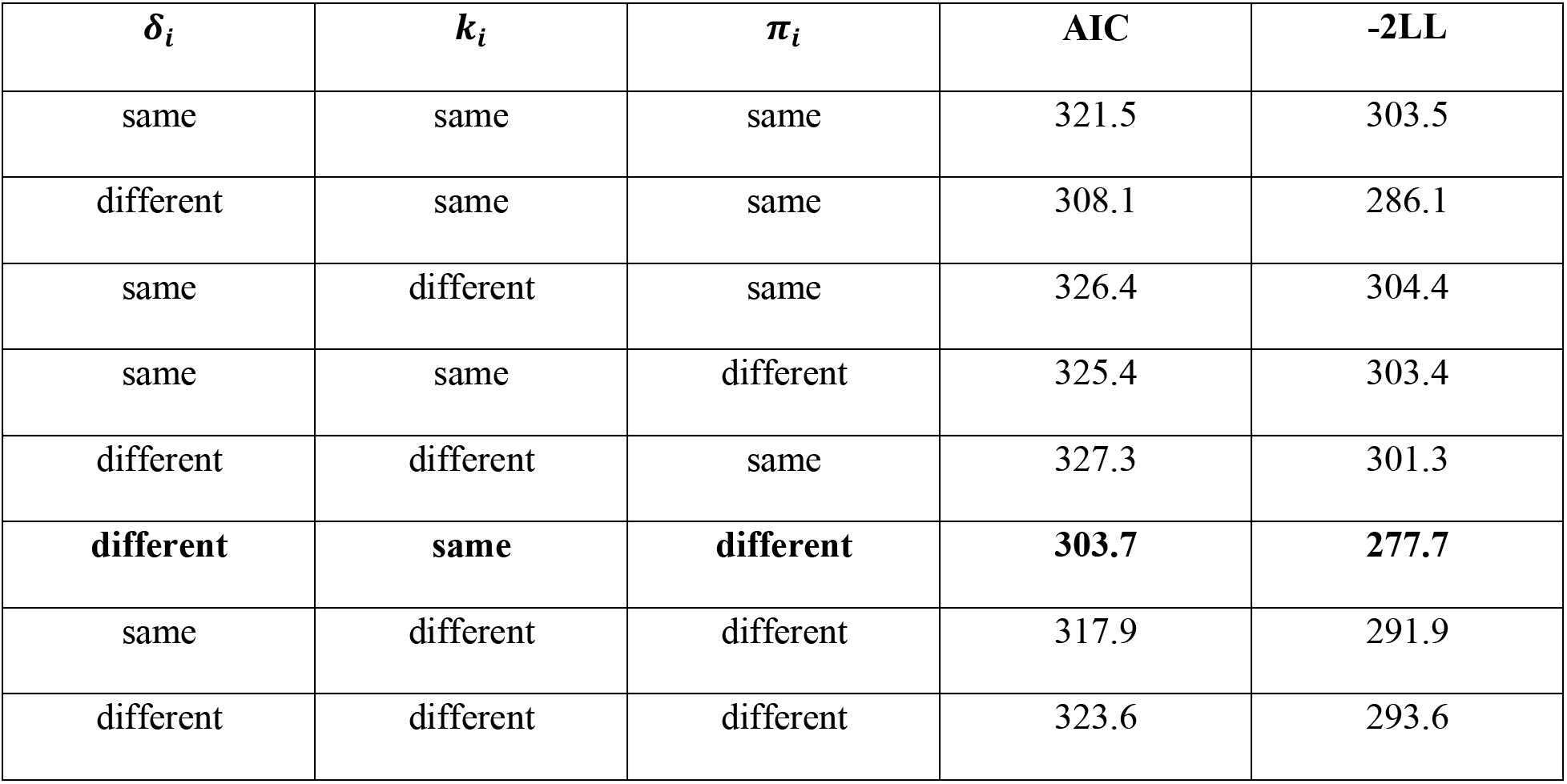
Different model structures explored while fitting nasal and BAL viral loads in only untreated animals. The model with the lowest Akaike information criteria (AIC) is best supported by the data (denoted in bold). ‘Same’ implies that the parameter takes the value from the same distribution in two spatial compartments whereas ‘different’ implies that the parameter has different distributions in two compartments. Here, -2LL represent −2 times the log-likelihood.

| $\delta_i$ | $k_i$ | $\pi_i$ | AIC | -2LL |
| --- | --- | --- | --- | --- |
| same | same | same | 321.5 | 303.5 |
| different | same | same | 308.1 | 286.1 |
| same | different | same | 326.4 | 304.4 |
| same | same | different | 325.4 | 303.4 |
| different | different | same | 327.3 | 301.3 |
| **different** | **same** | **different** | **303.7** | **277.7** |
| same | different | different | 317.9 | 291.9 |
| different | different | different | 323.6 | 293.6 |

### Model fit to viral load data from untreated rhesus macaques

The best model fit recapitulated the frequently observed trend of higher viral loads in BAL at late timepoints relative to the nasal passage in untreated macaques **(Fig 5a)**. The model also closely captured the viral dynamics in BAL from untreated animals and mostly captured nasal viral loads as well, though outlier datapoints compromised model fit somewhat in several of the animals.

**Figure 5:**
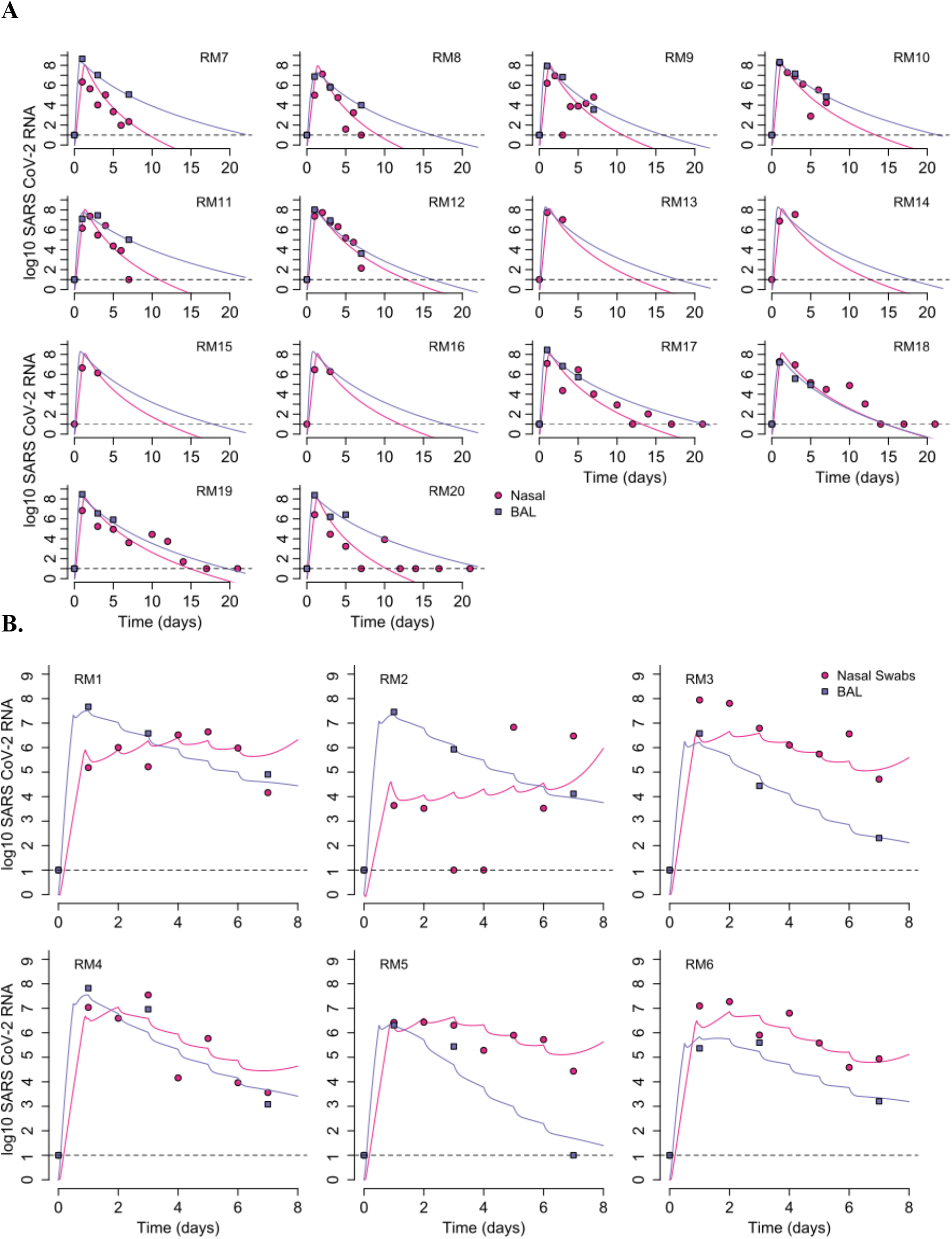
Mathematical model fits to viral load data. **A.** Fits to untreated animals. RM7-RM12 received a placebo vehicle in direct comparison to the remdesivir treated animals. RM13-20 are from different studies. **B.** Fits to 6 treated animals who received 10 mg/kg at day 0.5 and 5 mg/kg at days 1, 2, 3, 4, 5 and 6. Dots (pink=nasal swabs, purple = BAL) are datapoints and lines are model projections. Dots overlying the dotted line are below the limit of detection. Time is in days from infection.

Infected cell death rates were generally higher while viral replication rates were uniformly lower in the nasal passages relative to lungs in the untreated animals **(Table 4)**. The density-dependent exponent had a similar value in both compartments (*k*=0.09), was similar to that predicted in humans,^8^ and led to a 2-2.5-fold increase in the overall death rate of infected cells at peak viral load.

### Model fit to viral load data from remdesivir treated rhesus macaques

The PK/PD model (the combination of the PK model output and the viral dynamics model shown in Fig 4b) recapitulated the observed trend of lower viral loads in BAL at late timepoints relative to the nasal passages **(Fig 5b)**. The model also captured BAL viral loads on treatment accurately while reproducing nasal viral loads as well, though outlier datapoints again compromised model fit somewhat in several of the animals. The waning of drug levels and potency after dosing is evident in simulated tracings that exhibit decelerating viral decay rates after each dose.

### Increased remdesivir antiviral potency in nasal versus lung cells

The degree to which remdesivir suppressed viral replication varied somewhat across animals. Over the course of the treatment (from day 0.5 to day 7), the mean efficacy of the RDV treatment in nasal mucosa was estimated to be 88.4%, 91.4%, 86.9%, 81.0%, 86.2% and 83.5% in RM1, RM2, RM3, RM4, RM5 and RM6, respectively. Similarly, over the course of the treatment (from day 0 to day 7), the mean efficacy of the RDV treatment in lung was estimated to be 76.3%, 75.0%, 80.4%, 69.8%, 77.4% and 75.5% in RM1, RM2, RM3, RM4, RM5 and RM6, respectively **(Fig 6)**. Brief reductions in RDV drug concentrations between doses related to lower active metabolite levels in cells were associated with viral re-expansion after each dose **(Fig 5b)**.

**Figure 6:**
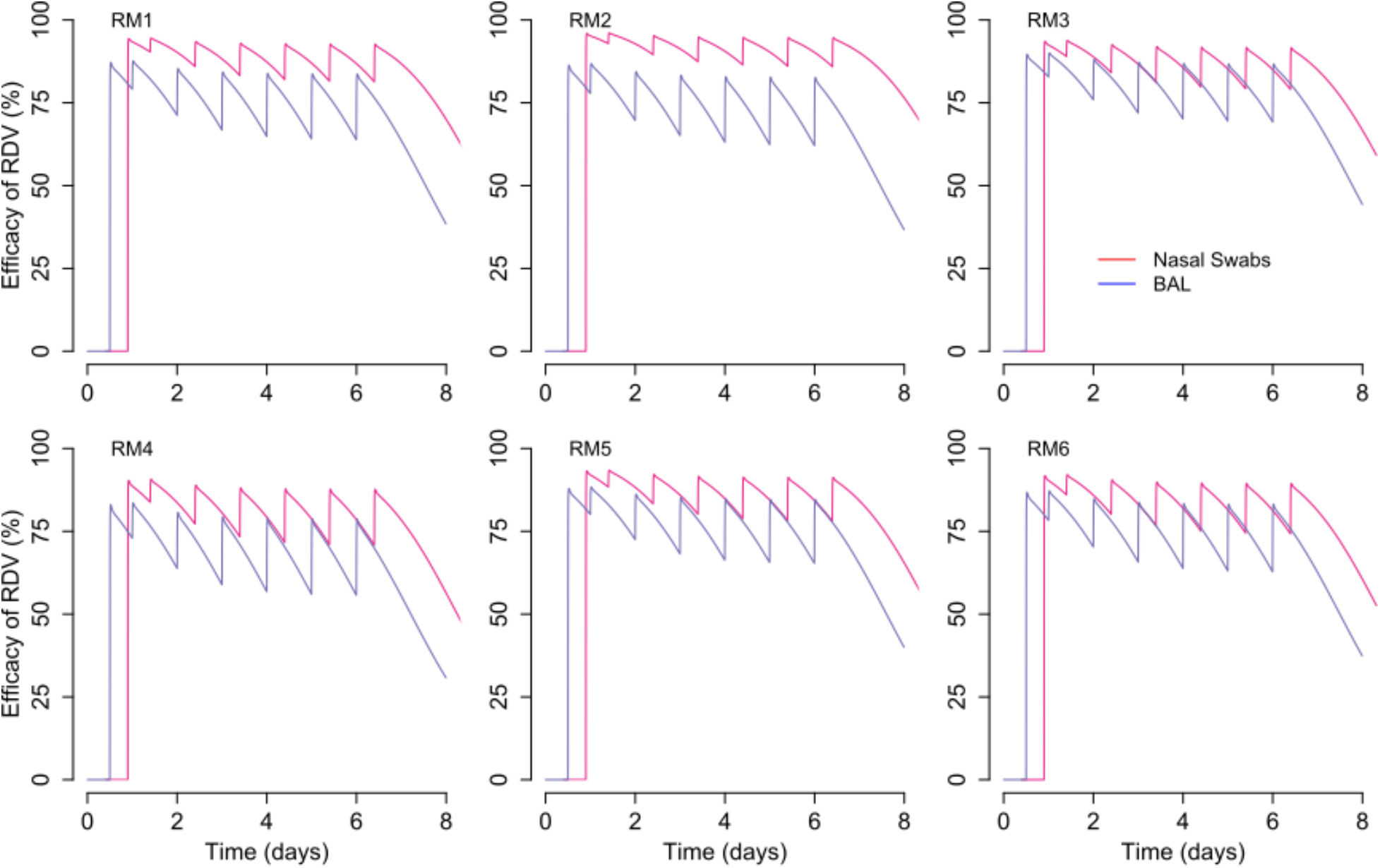
Projected direct antiviral efficacy of RDV treatment (∈_U_ and ∈_L_) in nasal passages (pink line) and the lung (blue line). Over the course of the treatment (from day 0.5 to day 7) the projected efficacy of remdesivir in nasal swabs (pink) is higher than in the lung (purple). Projections are based on data from RM1-6. Time is in days from infection.

The antiviral potency of remdesivir was estimated using “*in vivo*” EC_50_. Whereas “in vitro” IC50 estimate the antiviral concentration needed to inhibit 50% of viral replication based on *in vitro* experiments, we estimate EC_50_ based on viral loads measured *in vivo* in animal or human experiments.^10^ Estimates for *in ivio* EC_50_ were roughly 2-fold higher (2-fold lower potency) in the lung relative to the nasal passages **(Table 4)**. Variability in nasal viral load peak and contemporaneous viral loads between treated animals generally related to difference in *in vivo* EC_50_ rather than viral replication rate. RM2 had complex kinetics with low peak viral load followed by viral rebound (which was not captured by the model and may represent a drug resistant variant)^11^ and was found to have the lowest EC_50_. One animal with accelerated viral elimination in the lung (RM5) was found to have a higher infected cell death rate in lung but similar EC_50_ relative to the other 5 treated animals **(Table 4).**

### Lack of viral rebound in the lung is explained by infection dependent generation of refractory cells

We next performed counterfactual simulations in which the six treated animals were assumed to have not received treatment (ε_U_ =0 and ε_L_=0). The viral load trajectories in these simulations **(Fig. 7)** appear similar to those in untreated animals with BAL viral loads often exceeding nasal viral loads at later timepoints **(Fig 5a)**. Comparisons of the counterfactual viral load tracings to the treated animals suggests that a majority of viral load decrease in lungs is achieved following the first dose and is then carried forward throughout the duration of therapy with unchanged decay slopes. On the other hand, in nasal passages, viral load is decreased initially during therapy but then stabilizes or even increases, leading to higher viral loads than counterfactual projections **(Fig 7).**

**Figure 7:**
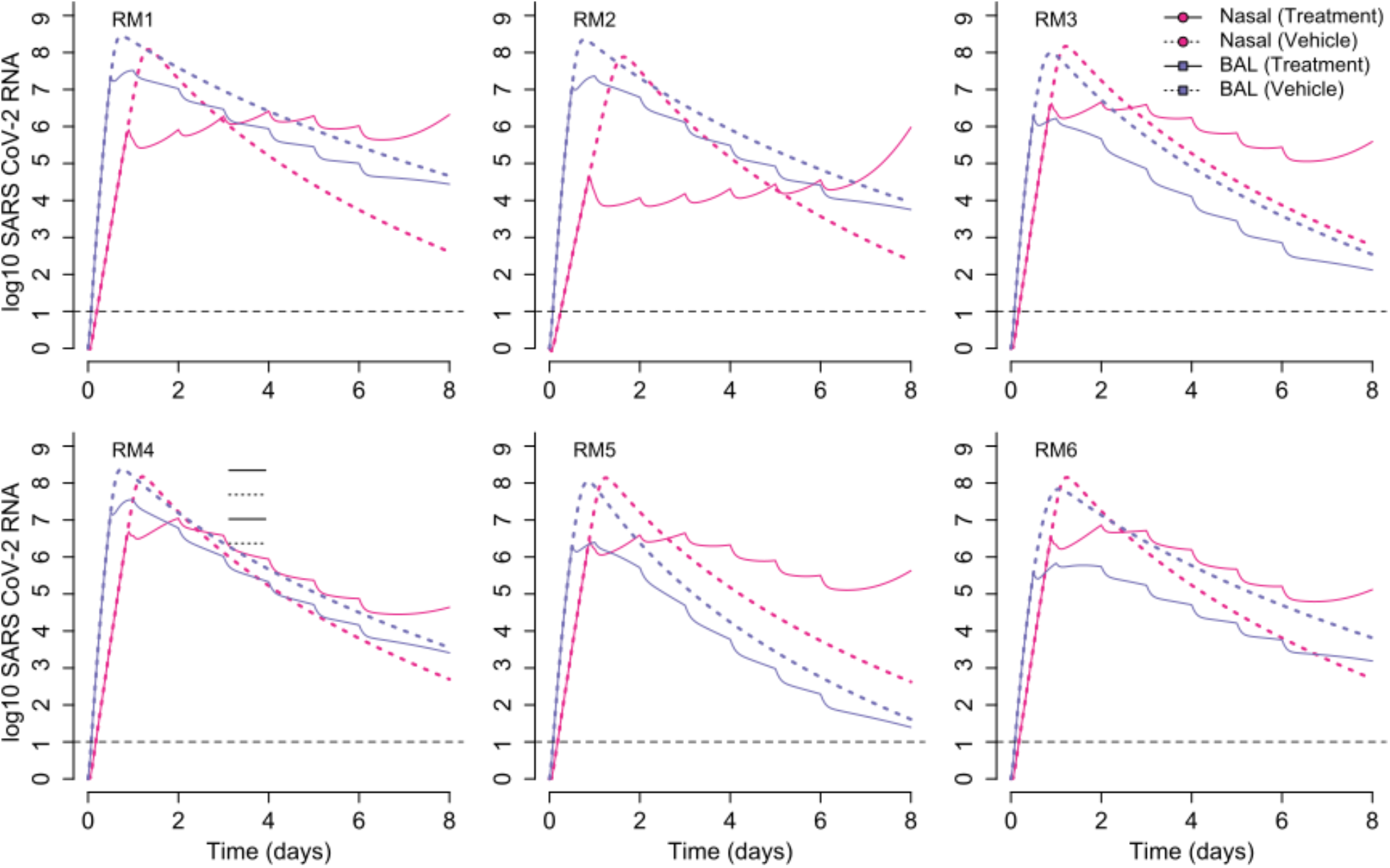
Projected impact of RDV treatment on viral dynamics in the nasal passages and lungs. Solid lines refer to the simulated viral loads under treatment, and dotted lines are counterfactual simulations assuming no treatment. In the case of the lung (BAL specimens), therapy is projected to lead to consistently lower viral loads. In the case of nasal viral load, therapy temporarily lowers viral load, but viral load is predicted to ultimately persist at higher levels than in the absence of treatment. Simulations are based on data from RM1-6. Time is in days from infection.

In the nasal cavity, somewhere between day 2 and 6 of therapy, the tracings cross and viral loads of the treated animals are predicted to exceed the counterfactual simulations of the same animals off therapy **(Fig. 7).** The model projects that early treatment reduces viral load, thereby decreasing new early infection and preventing depletion of susceptible cells in the nasal passages **(Fig 8)**. Even without assuming susceptible cell proliferation, there is an adequate population of these cells to establish a steady state of viral replication **(Fig 8).** In the lung where remdesivir is less potent and initially susceptible cells can become refractory to infection, treatment leads to a slower depletion of susceptible cells. These cells are nevertheless depleted in a linear fashion as they convert to a refractory state **(Fig 8)**. Inclusion of a refractory cell compartment is therefore necessary to allow linear elimination of virus from serial BAL samples.

**Figure 8:**
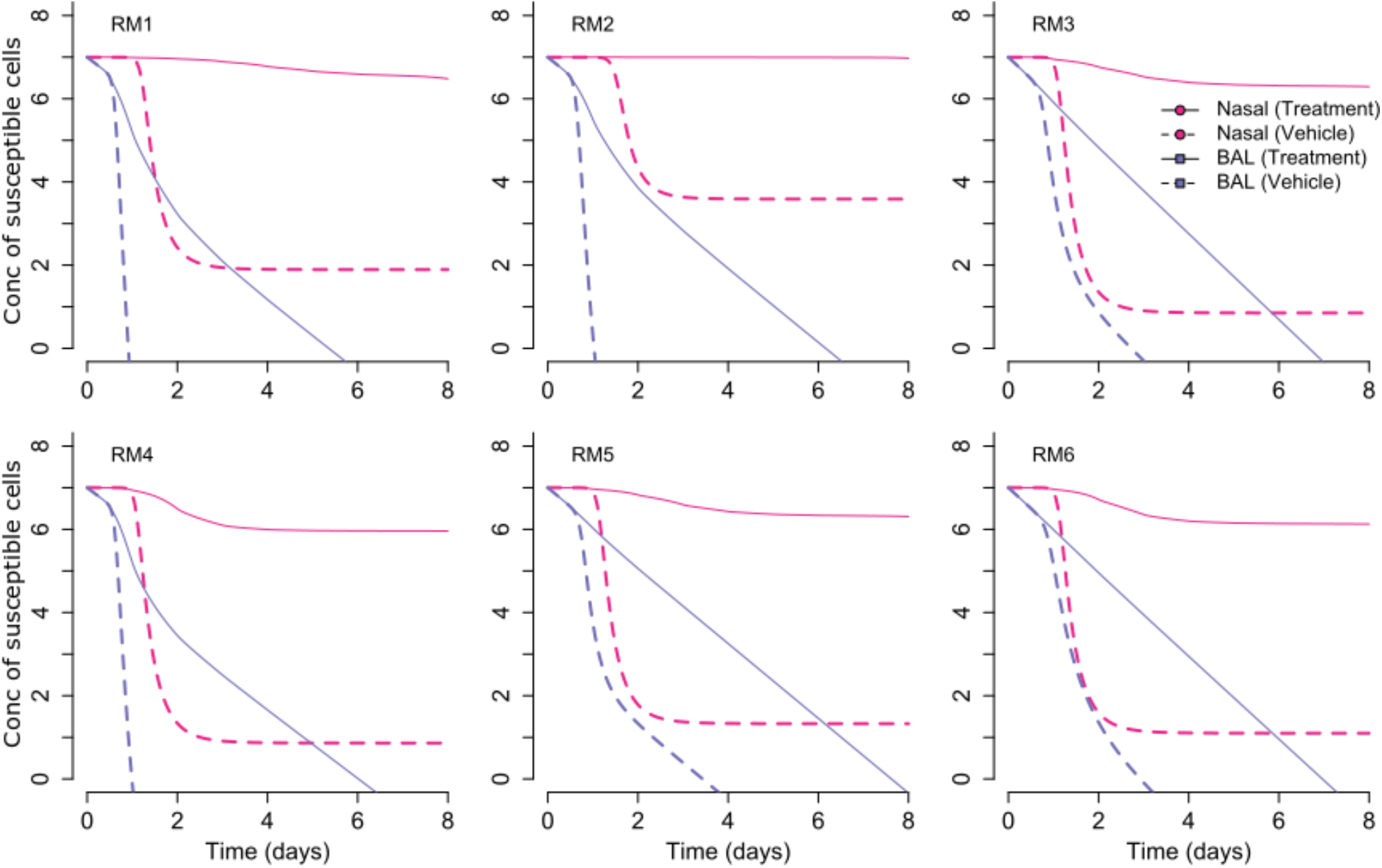
Mechanisms of lung protection in remdesivir treated animals. The concentration of susceptible cells is projected for simulations fit to treatment data (solid lines) and counterfactual simulations without therapy (dashed lines). In nasal passages, therapy limits initial depletion of susceptible cells which allows for persistent viral replication rather than elimination. In the lung (BAL specimens), treatment efficacy is lower and susceptible cells can become refractory to infection. The depletion of susceptible cells prevents persistent shedding. Simulations are based on data from RM1-6. Time is in days from infection.

### Decreased cell death in the lungs of remdesivir treated animals

As an informal assessment of lung damage, we longitudinally assessed cell death over time in our counterfactual simulations. In each case, therapy decreased the degree of peak cell death by at least 33% **(Fig. 9).** While lung damage is multi-factorial during COVID-19, this finding is qualitatively compatible with the observation that early remdesivir spared these 6 animals from severe clinical disease and abnormal lung histopathology.

**Figure 9:**
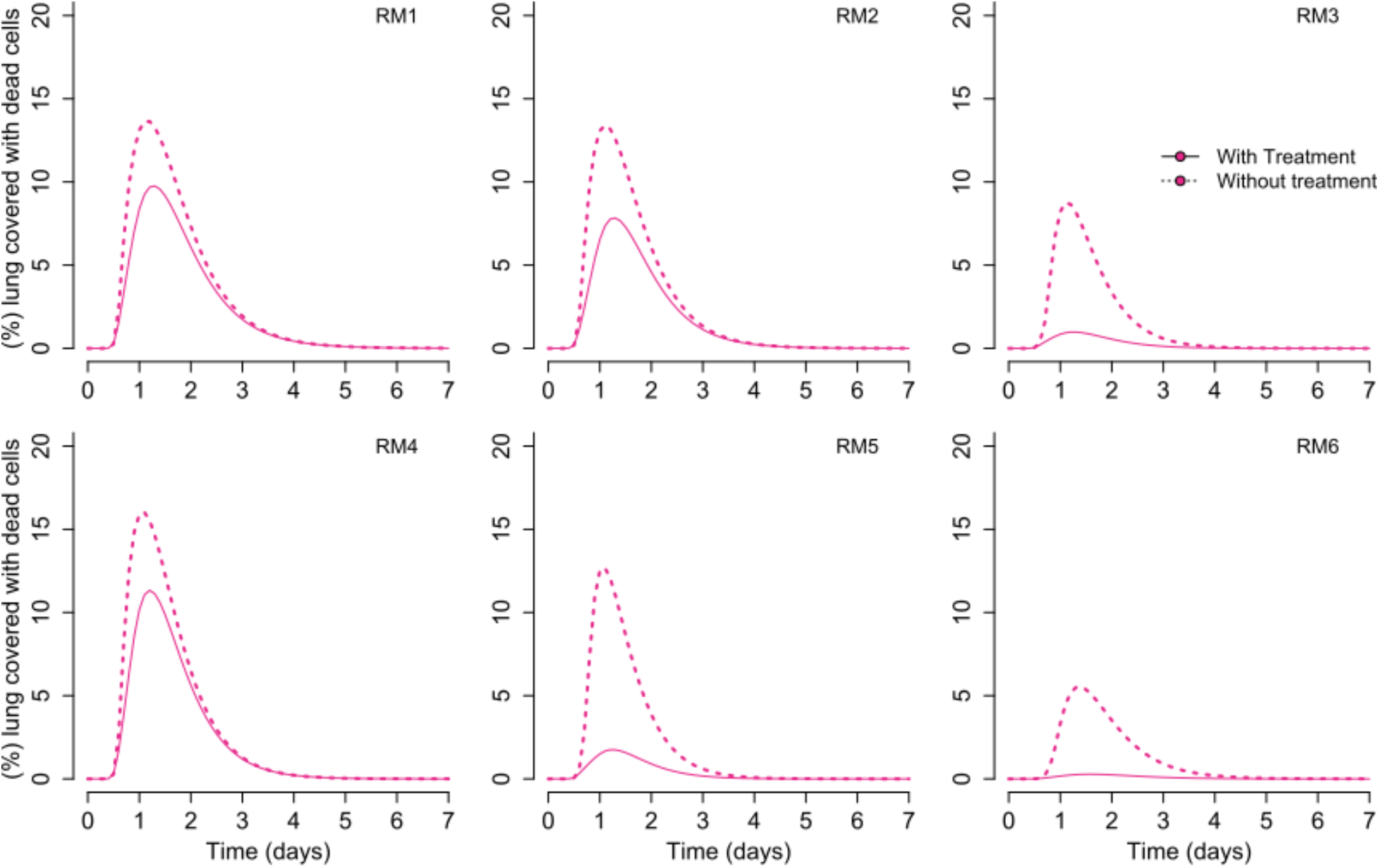
Simulated percentage of dead lung target cells as a proxy for lung damage. We define dead cells as the initial total of susceptible cells minus susceptible cells, infected cells and refractory cells in lung at each time point. Treatment lowers the percent of dead cells relative to the counterfactual simulations without therapy. Simulations are based on data from RM1-6. Time is in days from infection.

### Projected nasal and lung viral load trajectories at higher drug potency

Next, we performed sensitivity analyses in which we assumed a more potent antiviral effect, which could arise either from different dosing of remdesivir or a drug with a more potent drug. In nasal passages **(Fig 10a)** and in lungs **(Fig 10b)**, the impact of the first dose is more profound with higher potency leading to a more abrupt decline in viral load.

**Figure 10:**
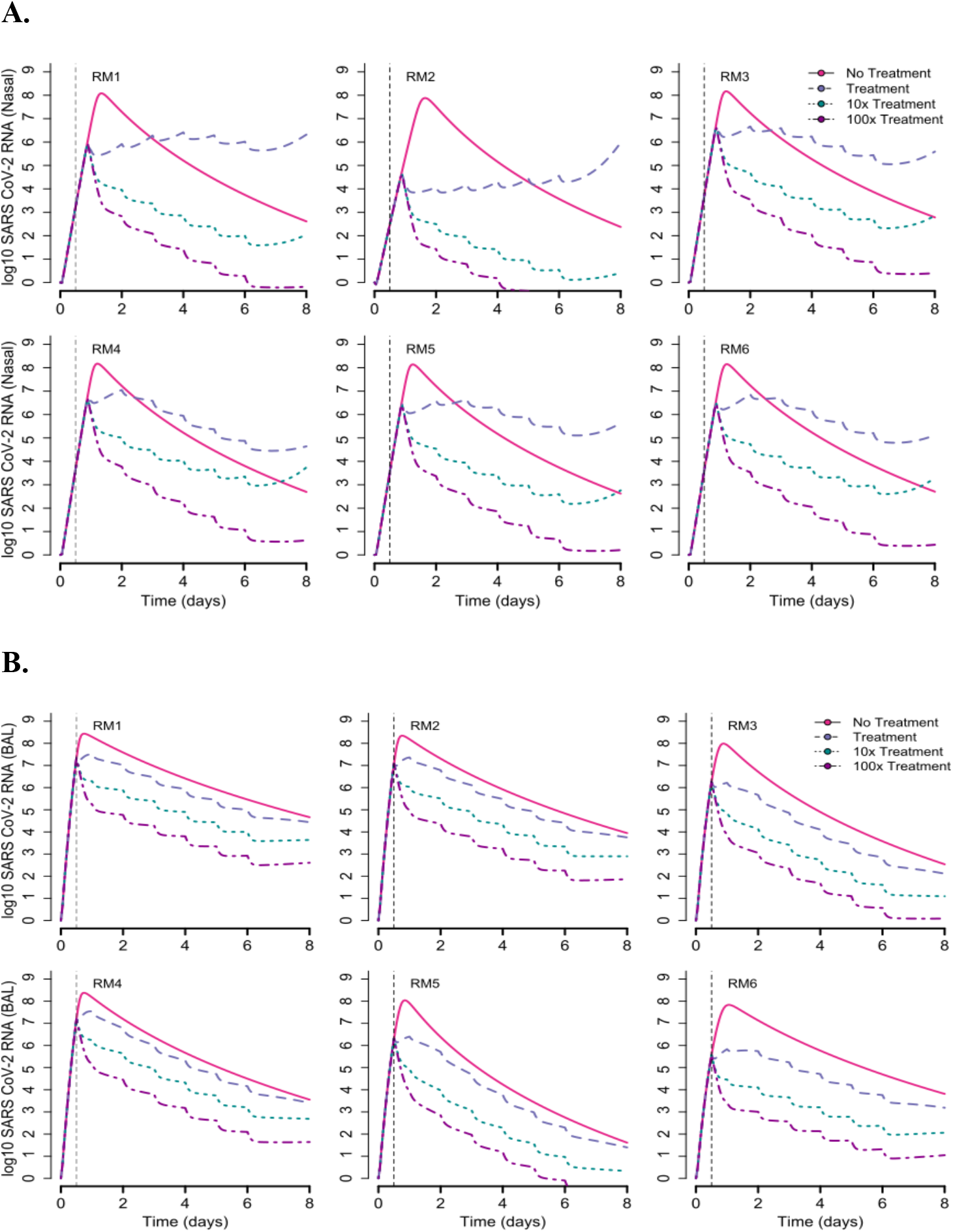
Predicted outcome of more potent remdesivir therapy. Therapy is simulated after lowering the *in vivo* EC_50_ 10-fold and 100-fold relative to data fitting in **Fig 5b.** Treatment is started 0.5 days after infection.**A.** Simulations of nasal viral load. **B.** Simulations of lung viral loads. Simulations are based on data from RM1-6.

We estimate that minimum drug efficacies of 99.99% and 99.5% would be required to eliminate virus from nasal passages and lungs within 5 days for a drug that is given 12 hours after infection. The need for such high potency reflects the lack of a concurrent immune response at this early stage of infection.

### Projected impact of later therapy in nasal and lung viral load kinetics

We previously predicted in modeling of human infection that antiviral treatment with moderate potency would not clear viral infection in the nasal passage (or sputum) if dosed prior to the peak viral load but would clear infection if dose several days later.^8^ Our simulations of the rhesus macaque data arrive at a similar conclusion in the nasal passage, that, paradoxically, later treatment with a moderate potency drug results in lower viral loads, whereas treatment started before peak results in increased late viral loads **(Fig 11a).** In contrast, in the lungs, later treatment at days 2 or 4 leads to a subsequent viral load trajectory similar to that of the earlier treated animals during the later stages of infection **(Fig 11b)**.

**Figure 11:**
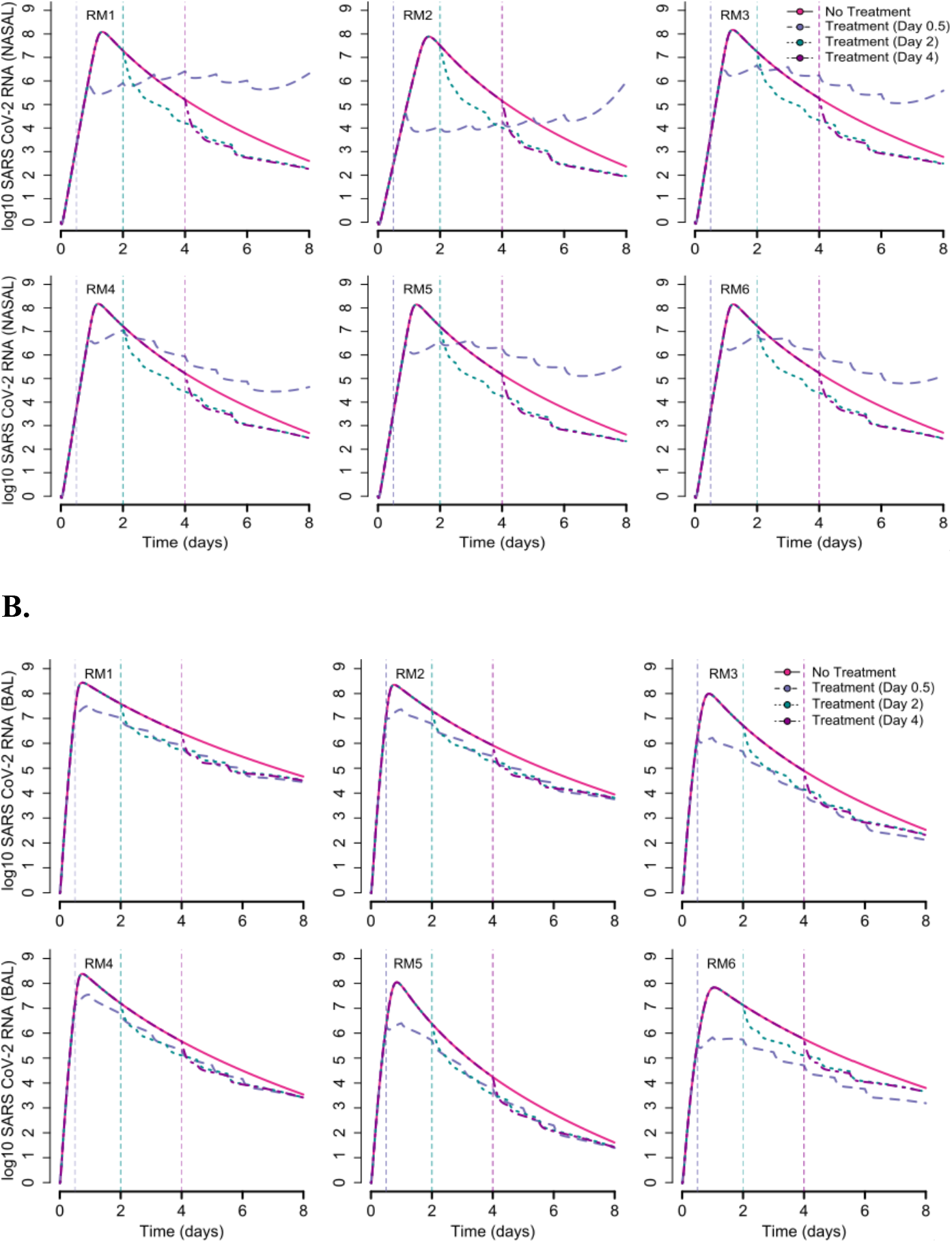
Predicted outcome of later remdesivir therapy. Therapy is simulated with the same antiviral potencies estimated from RM1-RM6 treated at day 0.5 but with initiation at later time points (days 2 and 4). **A.** Simulations of nasal viral load. **B.** Simulations of lung viral loads.

We estimate that minimum drug efficacies of 50% and 99.5% would be required to eliminate virus from nasal passages and lungs within 5 days for a drug that is given 4 days after infection. This result is due to the higher remaining viral load in the lungs of animals during the first untreated 5 days of infection.

## Discussion

Viral load is a valid surrogate endpoint for treatment efficacy of several viruses including HIV, hepatitis B, hepatitis C and cytomegalovirus.^12–16^ It is plausible that SARS CoV-2 lung viral load is also predictive of disease severity in humans. Viral loads from swabs of infected tissue provide an approximation of the number of infected cells at a given point in time, and therefore the surface area of infected tissue.^17,18^ Unfortunately, it is less certain whether viral load measurements can be leveraged for SARS CoV-2 treatment response in humans because BAL is required to measure lung viral loads but these are never performed longitudinally in infected people as part of routine clinical care. Experience from other respiratory viruses suggests that viral load measures in the upper airway by nasal swab or saliva may or may not be representative of those in the lung.^19^

Here we apply mathematical models to remdesivir treatment data in rhesus macaques in which both lung and nasal viral load were measured. We identify that the relationship between lung and nasal viral load in the context of antiviral treatment is complex and dependent on the potency and timing of therapy. While remdesivir lowered viral loads in the lung over the 7 days following infection, viral loads in the nasal mucosa were only transiently lowered. Several days into treatment viral loads actually increased slightly and surpassed what might have occurred without treatment. This result suggests that nasal viral load may not be an optimal surrogate for lung disease in the context of a partially effective antiviral therapy such as remdesivir at the doses used in this study. On the other hand, when we assume a more potent therapy with a lower *in vivo* EC_50_, then nasal viral loads are predicted to decrease in a linear fashion, in lock step with lung viral loads, immediately after starting treatment. Therefore, nasal viral loads in humans, measured either by duration of shedding or viral decay slope, may be a viable surrogate endpoint for lung viral load and downstream lung damage, but only in the context of a highly potent agent.

The experimental results highlight inherent strengths and limitations of the rhesus macaque model. Nasal passage viral kinetics and histologic lung damage appear similar between humans and rhesus macaques.^8,20^ We are also encouraged by the fact that a nearly equivalent mathematical model with a similar parameter set explains nasal viral loads in humans and rhesus macaques during the first week of infection,^8^ (though the acquired immune response is not modeled in the macaques because we do not observe complete viral elimination within the experimental timeframe). Similarly, our modeling of human data led to the prediction that a semi-potent treatment given extremely early infection might allow higher late nasal viral loads,^8^ which was also observed in the rhesus macaque experiments described herein.

On the other hand, in rhesus macaques, extensive lung damage and clinic illness is observed within two days of infection, which is not in keeping with severe illness in humans which emerges at least a week after the initial phase of illness.^5,21^ We hypothesize that direct intratracheal inoculation of macaques with a high viral titer results in more immediate infection of lung. In humans, a more common pattern is for respiratory viruses to start replicating in the upper airway and then transmit to the lungs in a second stage of infection.^22^ An alternative, and not mutually exclusive explanation is that the degree of viral replication in the lung can also be established extremely early in humans, but that the more extensive immune-mediated damage which may be correlated with the extent of early viral replication, occurs 1-2 weeks later. Had the rhesus macaques with the highest lung viral loads been followed for a longer period of time, it is possible that a more severe pneumonia would have developed at later timepoints.

A counterintuitive result predicted by our model is that remdesivir is slightly more potent in the nasal cavity than in the lung on a per cell level (assuming that drug levels are indeed equivalent in the two compartments). Nevertheless, SARS CoV-2 is not cleared in nasal passages as effectively as in the lungs while on treatment, because the effectiveness of antiviral therapies is never independent of the concurrent intensity of the immune response to infection.^10,23^ We previously predicted that a more potent therapy is needed after 2 days of SARS CoV-2 infection relative to >5 days after infection because there is little innate immune pressure against the virus during its early expansion phase.^8^ As a result, despite a slight blunting of initial viral loads, virus will rebound or stabilize and end up at a higher viral level in the nose than in the absence of treatment.

Here, we recapitulate this finding in the nasal passages, but also predict why this does not occur in the lungs of macaques. In the lung as in the nasal cavity, we assume density dependent killing as a proxy for an intensifying innate response to a higher burden of infection. However, our model also suggests that ongoing infection drives a percentage of lung cells to become temporarily refractory to infection. Inclusion of this assumption is required to recapitulate lung viral load data and to explain the observation that lung damage is severely blunted in animals receiving treatment. This assumption is supported by modeling of influenza infection.^24^

There are several limitations of our approach. First, our approximation of lung damage is relatively coarse based on the complexity of this post-viral inflammatory process which may be mediated by factors other than number of infected cells. This is therefore a qualitative target of our modeling. Second, our fits to nasal viral load are imperfect which may be due to imprecision in viral load measurements as well as missed components within the model. In the case of RM2, there is substantial viral rebound that may be due to incomplete innate responses to the first pulse of infection, or to *de* novo drug resistance. Third, we only model early infection and therefore neglect the critical impact of the late acquired immune response.^25–27^

In conclusion, we demonstrate that in rhesus macaques, the non-linear forces governing SARS CoV-2 viral load trajectories in the lung and nasal passages differ substantially in the presence of a partially effective antiviral therapy. To the extent that the rhesus macaque model approximates human infection, nasal viral load remains a promising surrogate endpoint marker, but perhaps only in the context of a highly potent antiviral therapy.

## Methods

### Experimental data

We analyzed viral load observations from nasal passages and BAL, and remdesivir and its metabolites plasma concentrations from 12 SARS-CoV-2-infected rhesus macaques in which 6 were treated with remdesivir and 6 received a vehicle control.^5^ Remdesivir was infused 12 hours after infection at a dose 10mg/kg with subsequent daily doses of 5 mg/kg for 6 day., We also added viral loads from nasal passages and BAL from 8 additional untreated animals from Muster et al.^7,21^ In both studies, rhesus macaques were infected with 2.6×10^6^ TCID50 of SARS-CoV-2 strain. Details about the infection and treatment protocol can be found in these two pre-prints.

We also analyzed more frequently sampled observations of remdesivir and its metabolites averaged from three healthy animals after a single IV infusion of 10mg/kg of remdesivir.^28^

#### Remdesivir pharmakinetics model

We used a compartmental and metabolism pharmacokinetics (PK) model for remdesivir. The goal of this model was to recapitulate the sparse data from remdesivir and its metabolites after several doses to the SARS-CoV-2-infected animals,^5^ along with the very frequently sampled data after a single dose in healthy animals.^6^ The PK model (depicted in Fig 2) describes the metabolism of remdesivir Prodrug GS-5734 (*A*_1_), to the alanine metabolite GS-704277 (*A*_2_) and subsequent parent Nucleoside GS-441524 (*A*_3_) in serum and their distribution to other tissue (*A*_1T_, *A*_2T_, and *A*_3T_ in the same order). Metabolism rates from GS-5734 to GS-704277 and to GS-441524 are describe by parameters *k*_12_ and *k*_23_. Drug distribution to other tissues and back to plasma are described by parameters *k*_1T_, *k*_1e_, *k*_2T_, *k*_2e_, *k*_3T_ and *k*_3e_. We assumed that in other tissues the active triphosphate metabolite (*A*_4T_) is metabolized from the parent nucleoside at rate *k*_34_. We finally assumed all metabolites have linear clearance with rates *k*_c1_, *k*_c2_, *k*_c3_, and *k*_c4_. These assumptions are captured by the differential equations below:

**Plasma compartments:**

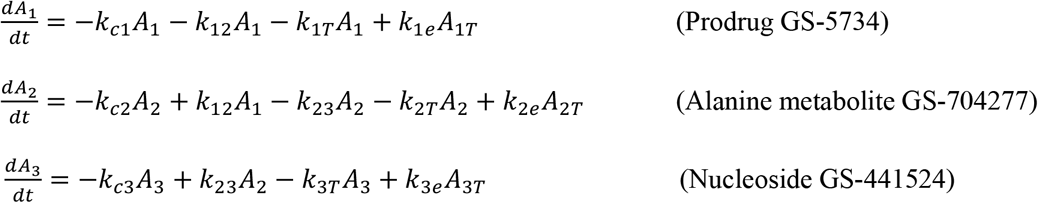

**Other tissue Compartments:**

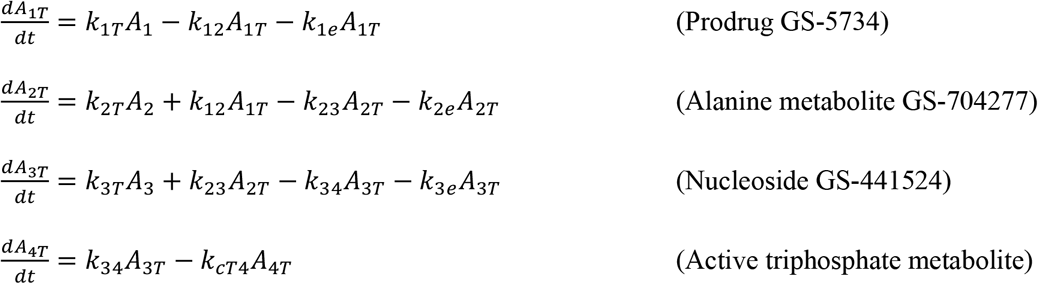

We fixed the half-life of prodrug GS-5734 in blood (ln 2 /*k*_*C*1_) to 1 hour, the half-life of alanine metabolite (GS-704277) in blood (ln 2 /*k*_*C*2_) to 24 hours,^28^ and the half-life of the active triphosphate metabolite (ln 2 /*k_cT4_*) to 24 hours.^6,7,29^ We fit the model to the data and estimated the remaining parameters.

#### Viral dynamics model

We extended our previous model of SARS Co-V-2 dynamics,^8^ to include both the lung and nasal passages. In both compartments (*i* ∈ [*L, U*], *L* for lung and *U* for NASAL), we assume that susceptible cells (*S_i_*) are infected at rate *β_i_V_i_S_i_* by SARS-CoV-2 (*V_i_*). SARS CoV-2-infected cells (*I_i_*) die with density dependent rate 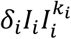, where *k_i_* describes by how much the first death rate depends on the infected cell density.^30^ This density dependent term represents a combined death of infected cells due to cytopathic effects of the virus and the killing of infected cells due to early immune responses. SARS CoV-2 is produced at a rate π_i_ and cleared with rate *γ*_i_.^24^ Free virus is exchanged between the lungs and nasal passages at rates *θ_LU_* and *θ_UL_*.

We also considered the possibility of the emergence of refractory cells. Due to antiviral actions of cytokines such as interferon, it has been experimentally demonstrated that uninfected lung airway cells may become refractory (*R_i_*) at rate *γ_i_*,^24^ and that infected cells may convert directly to refractory cells (*R_i_*) at rate *ϕ_i_*. Refractory cells may lose their refractory state and become susceptible at rate *ζ_i_*.^24^ Since we were only interested in the viral dynamics in a short span of ~7 days (with or without treatment), we ignored the death rate of uninfected and refractory cells in the lung, that are usually long-lived.

We also included the possibility of regeneration of susceptible cells during infection. Innate immune cells eliminate virus but can also induce pulmonary tissue damage or endothelium damage as part of this process.^31,32^ The restoration of the respiratory epithelial barrier after an injury is important and may happens within days after viral clearance,^33–35^ depending on the severity of the infection and the extent of lung involvement. Indeed, the proliferation of epithetical cells and progenitor stem cells (or, distal airway stem cells or DASCs) is critical for barrier repair following an inflammatory insult. Following lung injury, the tissue repair process is promoted by immune cells including innate lymphoid cells (ILC-IIs) and macrophages.^36^ Epithelial restoration is initiated locally by proliferating alveolar type II (AT2) cells.^37^ We modeled this restoration by adding a logistic proliferation of susceptible and refractory (but not infected^38^) epithetical cells with maximum rate *r_i_*. All the previous mechanisms are modeled by the following differential equation system:

**Nasal compartment:**

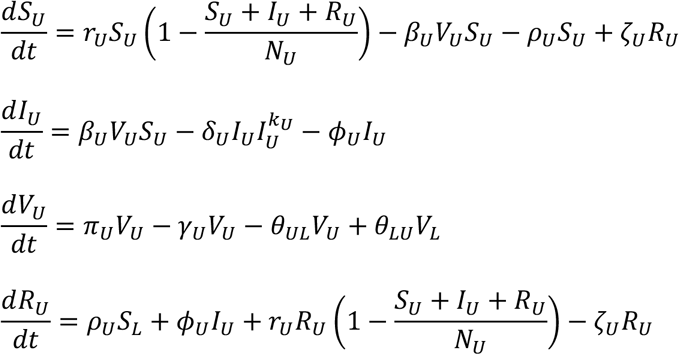

**Lung compartment (measured with BAL):**

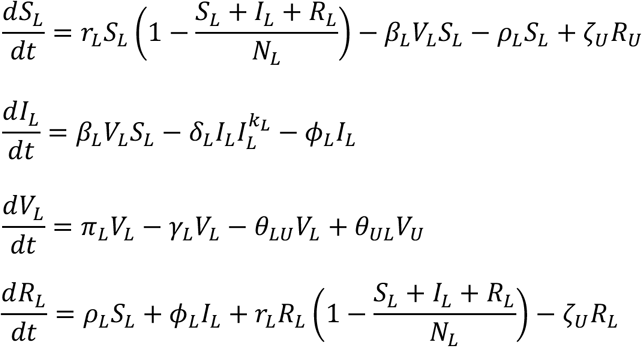

Here, *N_U_* and *N_L_* are the maximum carrying capacity of cells in respective compartments (assumed to be the total number of susceptible cells at time of infection).

#### Model assumptions about lung lesion formation

Although the formation of lung lesions during viral respiratory infections is multifactorial and complex we assumed it is mainly related to the number of dying SARS-CoV-2-infected cells and the lungs ability regenerate the epithelium damaged to avoid pulmonary edema.^33,39^ Thus, we modeled an informal surrogate for lesion damage (*G_L_*) with expansion kinetics equal to the total number of dying infected cells and shrinkage kinetics defined by the proliferation of susceptible and refractory cells representing the recovery of the lung tissue damage (or the reduction of the area covered by virus-induced lesions). *G_L_* = *N_L_* − *S_L_* − *I_L_* − *R_L_*

Taking derivative on both sides, we obtain

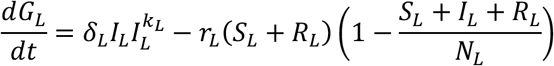

Notice that this definition of *G_L_* is equivalent to *G_L_* = *N_L_* − *S_L_* − *I_L_* − *R_L_*. Under this assumption, the fraction of the lung covered with dead cells would be: 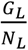.

#### Modeling remdesivir therapy

Here we assumed that RDV inhibits viral production. The effect of treatment on the viral production *π_i_* is reduced by a factor 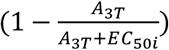, where *EC*_50*i*_ is the *in vivo* EC_50_ of the nucleoside GS-441524 in the respective compartment *i*.

#### Model fitting and selection

To fit different versions of the virus dynamics model to the data we used a non-linear mixed effects approach.^40,41^ Briefly, in this approach observed viral load for animal *k* at time *j* is modeled as 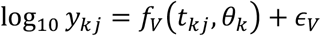 being *f_V_* the solution of model for the virus given the individual parameter vector *θ_k_* and *∈_V_* the measurement error. Here, the individual-parameter vector *θ_k_* is drawn from a population probability distribution. We estimated population parameters using the Stochastic Approximation Expectation Maximization (SAEM) algorithm and the individual parameters using a Markov Chain Monte-Carlo (MCMC) algorithm based on the estimated population distributions. Both algorithms, SAEM and MCMC, were performed using the software Monolix.

We first fit models to nasal and BAL viral loads from untreated animals assuming absent of cell proliferation and refractory cells. Given the lack of observations for the viral load upslope in BAL we assumed *β_L_* = *β_U_* (the heterogeneity in early viral dynamics in two regions can still be captured as it depends on the ratio 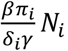). We excluded treated animals in this fitting procedure as *π_i_* and *EC*_50*i*_ cannot be estimated together. We assumed *t* = 0 as the time of infection with initial values *V_i_*(0) = 10 cps/ml, *I_i_*(0) = 0 cells/μl and *S_i_*(0) = 10^7^ cells/μl. We also assumed a virus clearance rate to be the same in both compartments *γ_L_* = *γ_U_* = 15/day and that *N_L_* = *N_U_* = 10^7^ cells/mL. We estimated the remaining parameters depending on each model assumptions. The explored competing models on this staged are listed in **Table 2**.

We next fit models to viral load and lung lesion observations from treated and untreated animals. Here, we explored different competing models listed in **Table 3** and described below. We explored models that included cell proliferation and refractory cells in the lungs, fixing *γ_U_* = 0,^42^ *ϕ_U_* = 0 and *ζ_U_* = 0. We explored the possibility that AT2 cells proliferate with maximum rate *r_L_* after some delay,^33^ i.e. *r_L_* = 0 if *t* < *τ*. We also included models assuming that the antiviral activity of remdesivir in nasal passages occurs after its activity in the lungs by a delay *v*. For comprehensiveness, we checked two models where refractory cells emerged from susceptible cells in a nonlinear fashion dependent on the concentration of infected cells: (1) with rate *ρ_i_S_i_I_i_* and (2) with rate 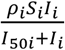. Notice that for the latter when *I*_50*L*_~0 then 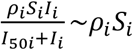. We finally checked if refractory cells could lose their refractory state and become susceptible cells. In all models we fixed *β* to the estimated value of the best model when fitting untreated animals and assumed remdesivir reduces virus production *π_i_* by a factor 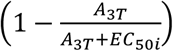 for each compartment *i*. Since we were estimating both *EC*_50*i*_ and *π_i_* together we explored fixing the standard deviation of the random effects of *EC*_50*i*_ to 0.1, 0.2 and 1. Finally, we only estimated the fixed effects of *r_L_, τ* and *v*, when applicable. Here, we also assumed *t* = 0 as the time of infection with same initial values and fixed parameters *γ_L_, γ_U_, N_L_* and *N_U_* as before. We estimated the remaining parameters depending on each model assumptions.

**Table 3:** Different model structures explored while fitting nasal and BAL viral loads in untreated and remdesivir treated animals. The model with the lowest Akaike information criteria (AIC) is best supported by the data (denoted in bold). Models with the inclusion of refractory cells and the proliferation of susceptible cells in lung but not in the nasal passage are better equipped to explain the reduced lung damage in treated animals.

| Refractory cells | Proliferation terms | Result |
| --- | --- | --- |
| No<br>($\rho_i's=0$ and $\phi_i's=0$) | No<br>($r_i's=0$) | AIC= 739.4 |
| Yes<br>($\rho_i's=0$ and $\phi_i's\neq 0$) | No<br>($r_i's=0$) | AIC=744.6 |
| Yes<br>($\rho_i's\neq 0$ and $\phi_i's=0$) | No<br>($r_i's=0$) | AIC=727.7 |
| No<br>($\rho_i's=0$ and $\phi_i's=0$) | Yes<br>($r_U=0$ and $r_L\neq 0$) | AIC=742.7 |
| Yes<br>($\rho_i's=0$ and $\phi_i's\neq 0$) | Yes<br>($r_U=0$ and $r_L\neq 0$) | AIC=730.3 |
| **Yes**<br>**($\rho_i's\neq 0$ and $\phi_i's=0$)** | **Yes**<br>**($r_U=0$ and $r_L\neq 0$)** | **AIC=722.4** |

To determine the best and most parsimonious model among the instances above, we computed the log-likelihood (log *L*) and the Akaike Information Criteria (AIC=-2log *L*+2*m*, where *m* is the number of parameters estimated). We assumed a model has similar support from the data if the difference between its AIC and the best model (lowest) AIC is less than two.^43^

**Table 4:** Estimated parameters under the model depicted in Figure 4b. We also estimated only the fixed effects of three parameters, *τ* = 0.42 days, *v* = 0.38 days and *r_L_* = 2.0/day while fixing *β* =1.7×10^-7^ virions^-^day^-1^.

| | $k_U = k_L$ | $\delta_U$ | Log10<br>($\pi_L$) | $\delta_L$ | Log10<br>($\pi_L$) | $\rho_L$ | $EC_{50U}$<br>(nM/gm) | $EC_{50L}$<br>(nM/gm) | CATEGOR<br>Y |
| --- | --- | --- | --- | --- | --- | --- | --- | --- | --- |
| RM1 | 0.09 | 0.91 | 2.5 | 0.51 | 3.2 | 1.9 | 0.8 | 1.9 | RDV |
| RM2 | 0.09 | 0.97 | 2.4 | 0.63 | 3.2 | 2.0 | 0.5 | 1.8 | RDV |
| RM3 | 0.09 | 0.85 | 2.6 | 0.96 | 3.2 | 2.4 | 0.9 | 1.5 | RDV |
| RM4 | 0.09 | 0.89 | 2.6 | 0.73 | 3.2 | 1.9 | 1.0 | 2.0 | RDV |
| RM5 | 0.09 | 0.89 | 2.6 | 1.26 | 3.1 | 2.1 | 0.8 | 1.6 | RDV |
| RM6 | 0.09 | 0.88 | 2.6 | 0.63 | 3.1 | 2.3 | 0.9 | 1.5 | RDV |
| RM7 | 0.09 | 1.14 | 2.5 | 0.51 | 3.2 | 1.9 | NA | NA | VEHICLE |
| RM8 | 0.09 | 1.20 | 2.5 | 0.66 | 3.1 | 2.8 | NA | NA | VEHICLE |
| RM9 | 0.09 | 0.99 | 2.5 | 0.71 | 3.2 | 2.0 | NA | NA | VEHICLE |
| RM10 | 0.09 | 0.80 | 2.6 | 0.53 | 3.2 | 2.0 | NA | NA | VEHICLE |
| RM11 | 0.09 | 0.99 | 2.5 | 0.46 | 3.1 | 2.5 | NA | NA | VEHICLE |
| RM12 | 0.09 | 0.83 | 2.6 | 0.71 | 3.2 | 2.0 | NA | NA | VEHICLE |
| RM13 | 0.09 | 0.85 | 2.6 | 0.64 | 3.2 | 2.0 | NA | NA | VEHICLE |
| RM14 | 0.09 | 0.83 | 2.5 | 0.64 | 3.2 | 2.1 | NA | NA | VEHICLE |
| RM15 | 0.09 | 0.89 | 2.5 | 0.64 | 3.2 | 2.0 | NA | NA | VEHICLE |
| RM16 | 0.09 | 0.89 | 2.5 | 0.64 | 3.2 | 2.0 | NA | NA | VEHICLE |
| RM17 | 0.09 | 0.80 | 2.5 | 0.53 | 3.2 | 2.0 | NA | NA | VEHICLE |
| RM18 | 0.09 | 0.71 | 2.6 | 0.76 | 3.1 | 2.7 | NA | NA | VEHICLE |
| RM19 | 0.09 | 0.70 | 2.5 | 0.58 | 3.2 | 2.0 | NA | NA | VEHICLE |
| RM20 | 0.09 | 1.05 | 2.5 | 0.48 | 3.2 | 2.1 | NA | NA | VEHICLE |
| **Median** | 0.09 | 0.89 | 2.6 | 0.68 | 3.2 | 2.0 | 0.8 | 1.7 | **RDV** |
| **Median** | 0.09 | 0.87 | 2.5 | 0.64 | 3.2 | 2.0 | NA | NA | **VEHICLE** |

